# Optimization of Functional Electric Stimulation for Foot Drop Patients using Inertial Measurement Unit

**DOI:** 10.64898/2026.06.14.732030

**Authors:** Muhammad Shahzaib, Usman Qamar Shaikh, Sadia Shakil, Sobia Janghsher

**Affiliations:** Biosignal Processing and Computational NeuroScience (BiCoNeS) Lab, Electrical Engineering Department, Institute of Space Technology, Islamabad, Pakistan; Turner Institute for Brain and Mental Health, Monash University, Melbourne, Australia

**Keywords:** Drop foot syndrome, Functional electrical stimulation, Gait phase detection, Optimization, IMU

## Abstract

Many people which are affected by drop foot syndrome, have to face difficulty while walking which leads to pathological gait. This type of syndrome is treated by means of an external artificial stimulation known as functional electric stimulator (FES). In this paper we are designing an online feedback control system which optimize the strength of a FES given to paretic muscle which results in correction of pathological gait of the patient in a tolerable domain. Different phases of gait are identified using inertial measurement unit (IMU) as a feedback sensor mounted on the foot. Data is collected form 8 different healthy subjects and average of collected data is used as a reference template. Different trajectories of drop foot patients are simulated (due to unavailability of patients) and corrected according to the reference template.

## I. Introduction

**S**TROKE is widespread in Pakistan and according to Pakistan Stroke Society (PSS) [1] stroke incidence in Pakistan is estimated to be 250 per 100,000, which estimated approximately 350,000 stroke patients every year in Pakistan. According to a recent report by WHO [2] stroke is the third leading cause of disability. 15 million people suffer from stroke every year, out of which 5 million are disabled permanently. The number of stroke incidences is doubled in low and middle income countries over the last 4 decades.According PSS the major cause of stroke is high blood-pressure and more than 30% of population above 45 suffer from high BP and large amount of population of strokes patients also suffers after accidents. As a result, the number of stroke patients in Pakistan is higher as compared to other developed countries of the world. The recovery for stroke patients, in general, is slow and mostly partial, which makes stroke the leading cause of disability worldwide. It is even a bigger issue in countries like Pakistan due to lack of resources as well as the number of rehabilitation centers. On average the daily number patients coming to rehabilitation centers are greater than 170 (according to NIRM rehab. Center Islamabad [3]). We don’t have any concrete policies at national/provincial levels to deal with such an alarmingly rising stroke-patients population. We have limited number of hospitals to take care of all the stroke survivors and most of them are not utilizing the multidisciplinary rehabilitation teams for these patients such as involving doctors, radiologists, and physiotherapists and state of the art technologies which are mostly expensive. Even the few places using these teams don’t take advantage of Electromyogram (EMG), Inertial measurement units (IMU’s) or other feedback sensors to collect data and utilize the results (feedback mechanisms) for rehabilitation purposes. Furthermore, due to lack of rehabilitation centers patients are mostly send home for rehabilitation. At home, the rehabilitation process may become slower or even stop due to many reasons, such as lack of caregiver’s relevant training, lack of resources to monitor the progress at home, and unavailability of the system to remotely communicate the progress with the doctors/hospitals and one of the most important parameter is the cost (roughly 5 to 10 lack Rs.) of feedback devices that assists the caregiver and analyze for rehab patients at home. This leads to a majority of stroke survivors not recovering to a level in which they can live a healthy life. People who recover from stroke often experience the inability to lift the foot. The condition is referred as drop foot syndrome (DF). DF patients are subjected to pathological gait. Due to that, their walking speed is significantly reduced and they drag the foot as they walk. The conventional solution to solve DF during gait is to wear mechanical ankle foot orthotics (AFOs). AFO limits the planter flexion that allows the foot to lift from ground during walk. However lack of muscle activity by using AFO can cause muscle wasting and joint stiffness. In most post stroke drop foot patients, the peripheral nervous system remains intact but the motor activity is lost. For that case artificial motor activity can be induced by using functional electrical stimulation technique. In human body the nervous system plays as a communication pathway between brain and other parts of the body. This nervous system further divided into two parts, central nervous system and peripheral nervous system (close to limbs). A normal human can contract their muscles using electric signals trasmitted by brain through nerves to the designated part of the body. These signals are known as Electromyography (EMG) signal. The basic building block of an EMG signal is called motor unit (MU) [4]. These motor units (MU) fire pulses due to the polarization and repolarization known as action potentials (AP) [4] in the muscle fiber membrane when contracted or active. Consequently, an EMG signal is the combination or overlapping of motor unit action potentials (MUAP) of different number of frequencies when fired continuously. If central nervous system is damage due to stroke or any spinal cord injury and peripheral nerves are still intact, then treatment of rehabilitation can be done by means of external assistive technology called Functional electric stimulation (FES). FES generate small electric pulses via skin using gel/dry electrodes non-invasively or by electrode implant, to contract desire muscles. These tiny signals activates the MU of muscle fiber membrane which in results contracts the muscles. The electrodes of FES could be implanted or attached to skin. The shape of the FES signal is a biphasic square wave (having net current zero) with an amplitude ranging from about 20 to 150 mA and pulse width of 10 to 500 µs. The frequency of FES ranges between 20 and 100 Hz. Due to the advancement of microcontrollers and compact sensors, a number of closed loop control system for neuroprosthesis are proposed in the 21st century however most validations relied either on simulations or sitting or lying subjects. For example a PID control system for neuroprosthesis is proposed by Benedict and Ruiz [5]. Similarly Hayashibe and Mitsuhiro [6] have suggested the use of predictive control. Both of the above researches were limited to simulated results. In addition to that, Valtin and Seel [7] performed a research and proposed iterative learning control and tested their algorithm on sitting subjects. Seel and werner [8] has developed a closed loop control system that corrects the pitch angle of foot by using iterative learning control. That algorithm is designed to adapt to stepping frequency. However the developed control system was validated only on 6 subjects with respectable results. The aim of this research is to develop a mathematical model of pitch angle during walk for physiological gait as well as pathological (DF) gait. by doing so, an optimized solution for drop foot patients will be proposed. This paper has six main sections; introduction, problem formulation, hardware designing, software and algorithm, results and conclusion. The rest of the paper is arranged in the following order, section I is introduction, section II contains problem formulation followed by the hardware designing in section III. The section IV provides the software and algorithm, and the section V is results and conclusion in section VI.

## II. PROBLEM FORMULATION

The block diagram of feedback optimization of FES intensity is shown in Fig. 1

**Fig. 1:**
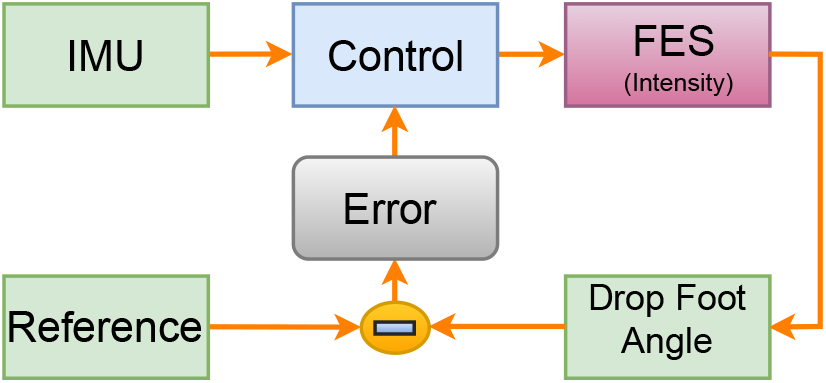
Block Diagram of FES Optimization

### A. Objective Function

The objective is to minimize the error between drop foot patient and healthy subject by controlling the intensity of FES in an optimal manner and is given by 1. Where HS represents healthy subject data or reference template, CT represents the corrected signal of drop foot patient after applying optimal FES intensity *α* and *x* is the time domain samples.

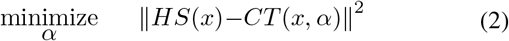

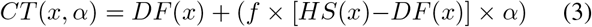

*DF* (*x*) is the time series model of drop foot patient and *f* is the factor that represents the variation in the signal of a drop foot patient. We have modeled the drop foot patient’s signal trajectory (as shown in Fig. 3) by applying regression on an artificial generated signal which is close to actual trajectory (Fig. 4 [9]) shown in Fig. 2 by red color. Since, this modeled signals slightly varies patient to patient, so a factor *f* will compensate this variation and its ranges lies between zero and one.

**Fig. 2:**
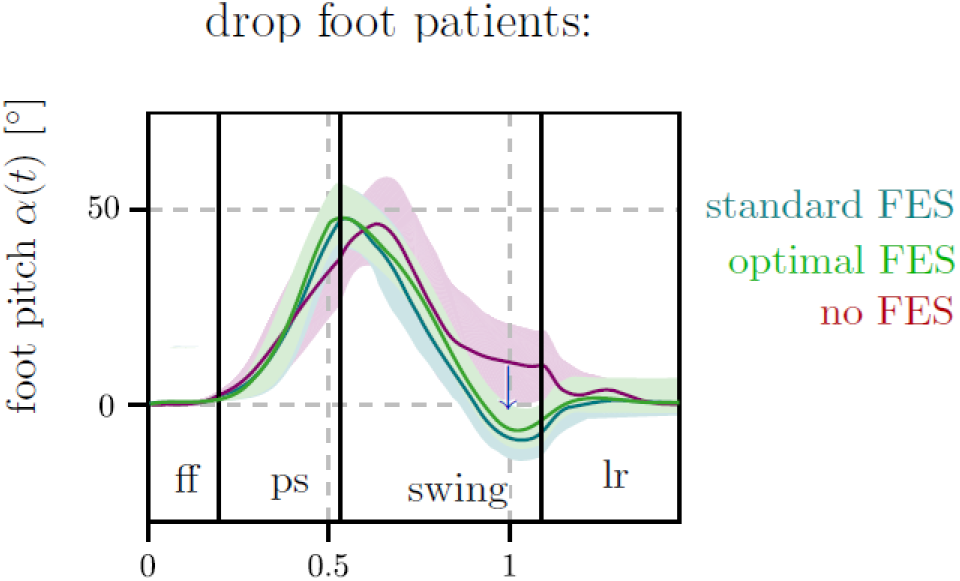
Drop foot patient signal [9]. Flat foot(ff), pre-swing (ps), lr (loading response)

**Fig. 3:**
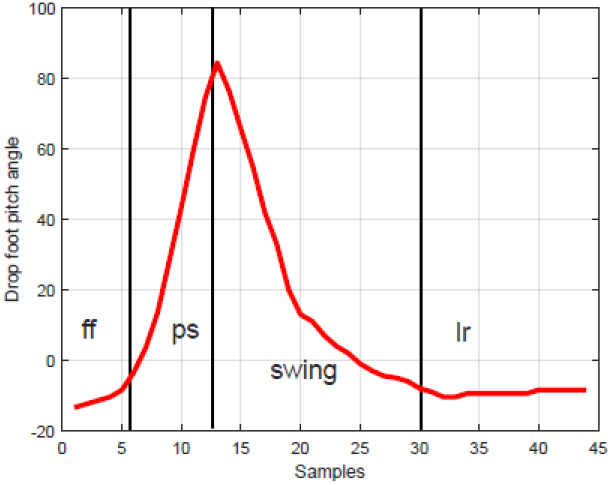
Drop foot patient generated signal

**Fig. 4:**
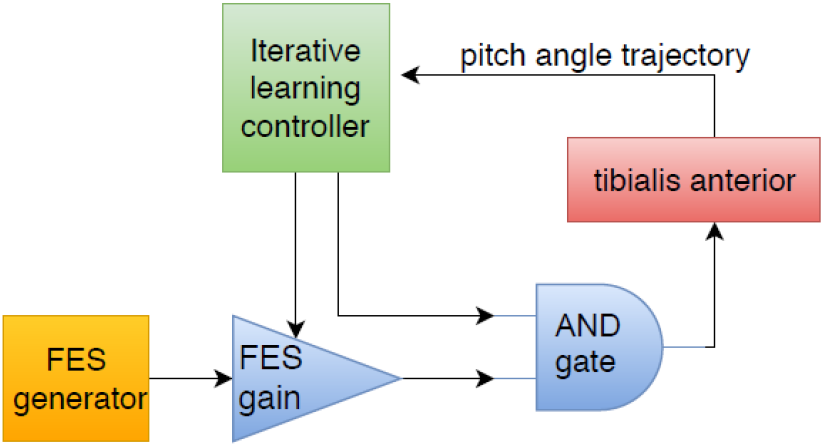
Drop foot patient generated signal

**Fig. 5:**
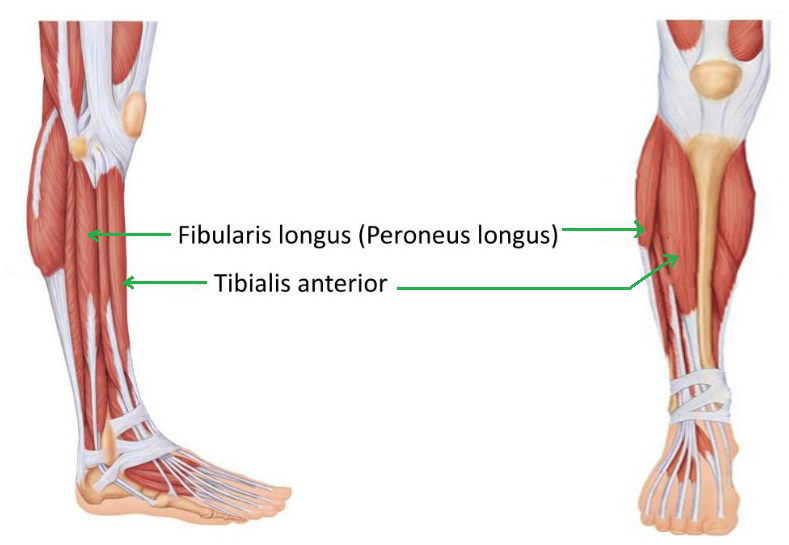
Paretic Muscles

**Fig. 6:**
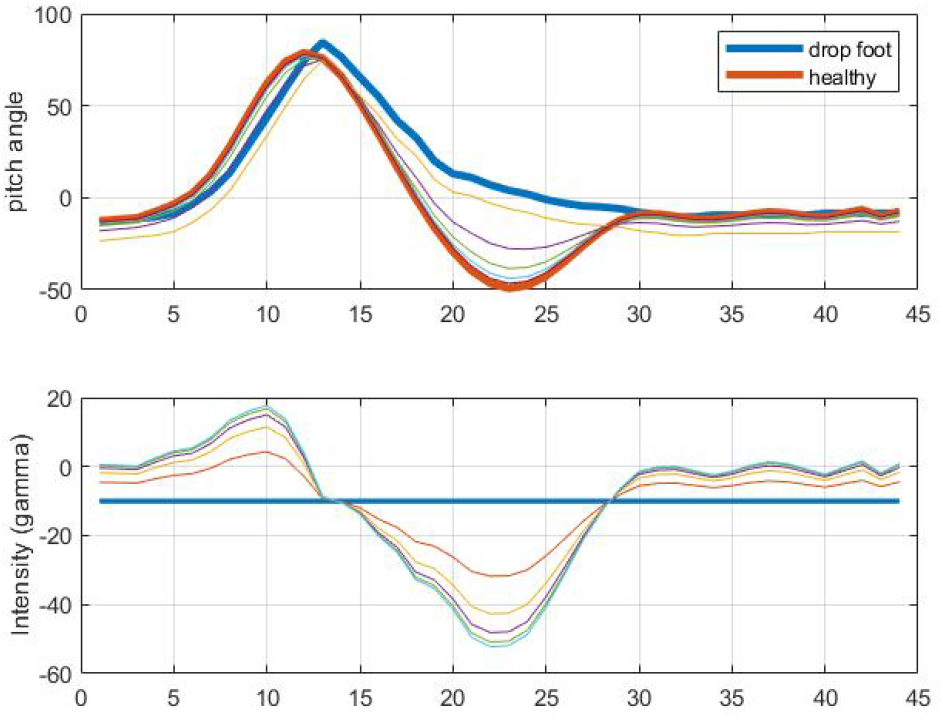
Drop Foot vs FES Values

**Fig. 7:**
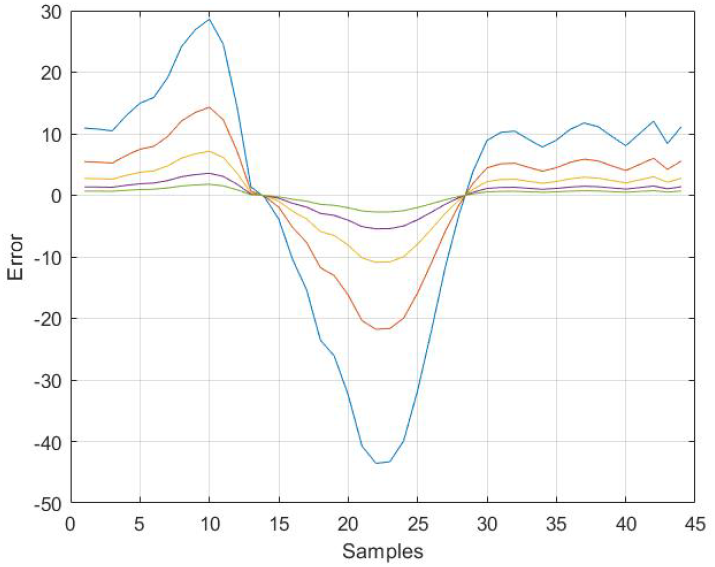
Error vs FES Values

**Fig. 8:**
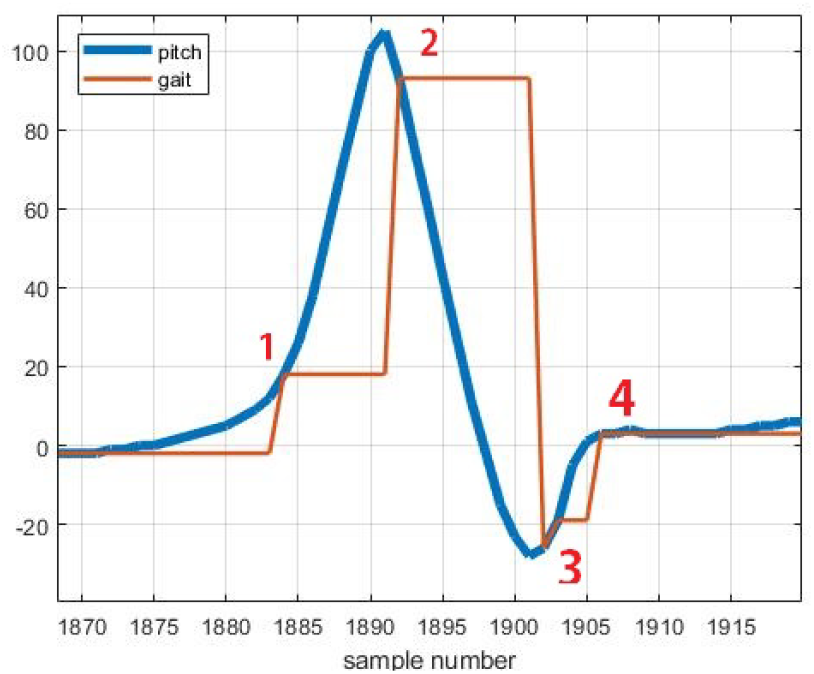
Gait Event detection. 1) Heel rise, 2) Toe off, 3) Heel strike, 4) Flat foot

### B. Constraints

This problem is an un-constraints problem and there is a boundary condition on the intensity (*α*) of a FES which lies in the range of 0 < *α* ≤ 50.

### C. Decision variable

The Intensity of FES (*α*) is the decision variable whose value determines the amount of error.

### D. Convexity

We check the convexity of a function, by taking double derivative of (3) w.r.t *α* we get zero. Secondly, we check the domain of a decision variable (*α*) that is also continuous. Since, our objective function is convex and domain is continuous so our problem is convex and Matlab CVX is used to solve this problem.

## III. SOFTWARE AND ALGORITHM

### A. CVX Implementation

A Matlab library of CVX is used to solve the problem (1), by generating a random number for factor f (i.e 0.7692) we get an error value of −1.152*e*^−8^ at *α* = 1.3. Since, we know the optimal value, we check response of the function around this optimal value of *α*.

### B. Iterative Learning Control system (ILCS)

1. Adaptation with walking speed: The FES need to be triggered 200 ms [8] before the toe off. This is because muscle contraction takes about 200 ms after the signal. During walk, there is significant difference in walking speed at every step. So the FES trigger time needs to be adaptive with walking speed. To achieve this, the time taken from heel rise to toe off is measured as Δt. The measure value indicates the walking speed or stepping frequency. The measured information is used to estimate the FES trigger time for the next step. The FES is triggered at some point between heel rise and toe off. The delay after heel rise to the point of firing FES is measured as follows: *Delay* = (Δ*t* − 100)*ms*. As soon as the heel rise in next step is detected, the future value Tf for FES trigger time is calculated as:

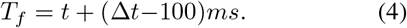
2. Closed loop control system for optimum FES strength: The main aim of this research is to develop a system that can provide optimum amount of FES charge to the patient. Excessive and unwanted charge can potentially damage the muscle fibers and can fatigue the muscles earlier. To overcome this problem, a closed loop control system is developed. The amplitude of FES charge is increased or decreased until the desired pitch is achieved. The amount of muscle contraction is proportional to the strength/charge of *FES*(*FES*_*c*_) and calculated as:

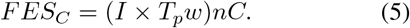

Where, *I* is the current and *T*_*p*_*w* is the pulse width in µs. The shape of FES pulse has a biphasic square wave shape in order to minimize charge accumulation in the muscle’s tissue and make the net current zero. A block diagram of feedback system is shown in Fig. 4. Desired pitch data from ten healthy subjects was gathered at different stepping frequencies. The desired pitch values during swing phase of foot is calculated.

## IV. BRIEF ANATOMY OF LOWER LIMB MUSCLES

The muscle responsible for foot lift is tibialis anterior which is innervated by sciatic nerve and fibularis longus muscle which is innervated by peroneal nerve is responsible for foot roll.

## V. RESULTS

### A. Gait event detection

The real time gait events are detected based upon the pitch velocity of foot. these events include flat foot start, heel rise, toe off and heel strike. the gait event detection is divided as follows. Flat foot is the duration when foot is completely on the ground. During this period, there is negligible change in pitch angle. So the pitch angular velocity is almost zero. Heel rise is the instance after the flat foot period. At this point of moment, heel is lifted and a sudden positive increase in pitch angular velocity is observed. The angular velocity remains positive until the toe off is achieved. After the heel rise, the final contact of foot with the ground is at toe. As soon as the toe is lifted from the ground, the pitch angular velocity changes sign. The foot remains in the swing phase till the heel strike. The swing phase of the foot ends at heel strike. During the swing phase the pitch angular velocity of foot remains negative. At the point of heel strike, the angular velocity changes sign. After the heel strike, the period during which foot accepts the weight of the body from opposite foot is called loading response. The start of the flat foot is nearly synchronous with the opposite heel rise. As the foot is being loaded to the ground after heel strike, a positive pitch angular velocity is observed for a short duration.

### B. FES pass length

The goal of the research is to facilitate the drop foot patient to lift foot during walk. Thus, FES only need to be applied during swing phase. The swing phase cover about 40 percent of the gait cycle. It started from toe off and ends at heel strike or initial contact. A problem arises that the muscle contraction after applying FES signal takes about 100ms. So the FES should be triggered 100 ms before the toe off. The gait event detection algorithm for predicting the toe off event before it occurs, is discussed in Section III-B. As soon as the heel strike is detected, the FES is stopped. In case of wrongly gait event detection, there is a check to stop the FES signal if time exceeds 2s.

## VI. CONCLUSION

In this research an adaptive optimized assistive device for drop foot patients is proposed to correct the pathological gait. The data is collected from 10 healthy subjects and an algorithm is modeled. The results are simulated in Matlab that clearly shows the transformation from pathological gait to physiological gate.

## VII. Acknowledgment

